# Contexts facilitate dynamic value encoding in the mesolimbic dopamine system

**DOI:** 10.1101/2023.11.05.565687

**Authors:** Kurt M. Fraser, Val L. Collins, Amy R. Wolff, David J. Ottenheimer, Kaisa N. Bornhoft, Fiona Pat, Bridget J. Chen, Patricia H Janak, Benjamin T. Saunders

## Abstract

Adaptive behavior in a dynamic environment often requires rapid revaluation of stimuli that deviates from well-learned associations. The divergence between stable value-encoding and appropriate behavioral output remains a critical test to theories of dopamine’s function in learning, motivation, and motor control. Yet how dopamine neurons are involved in the revaluation of cues when the world changes to alter our behavior remains unclear. Here we make use of pharmacology, in vivo electrophysiology, fiber photometry, and optogenetics to resolve the contributions of the mesolimbic dopamine system to the dynamic reorganization of reward-seeking. Male and female rats were trained to discriminate when a conditioned stimulus would be followed by sucrose reward by exploiting the prior, non-overlapping presentation of a separate discrete cue - an occasion setter. Only when the occasion setter’s presentation preceded the conditioned stimulus did the conditioned stimulus predict sucrose delivery. As a result, in this task we were able to dissociate the average value of the conditioned stimulus from its immediate expected value on a trial-to-trial basis. Both the activity of ventral tegmental area dopamine neurons and dopamine signaling in the nucleus accumbens were essential for rats to successfully update behavioral responding in response to the occasion setter. Moreover, dopamine release in the nucleus accumbens following the conditioned stimulus only occurred when the occasion setter indicated it would predict reward. Downstream of dopamine release, we found that single neurons in the nucleus accumbens dynamically tracked the value of the conditioned stimulus. Together these results reveal a novel mechanism within the mesolimbic dopamine system for the rapid revaluation of motivation.

## INTRODUCTION

The ability to determine the importance of environmental stimuli and generate appropriate behavioral responses is essential for survival. Dopamine neurons in the midbrain are thought to be critically involved in one aspect of this evaluative process – determining the expected value of the outcome associated with a given stimulus (Rescorla and Wagner, 1972; Schultz et al., 1997; Schultz and Dickinson, 2000; Fiorillo et al., 2003; Pan et al., 2005; Tobler et al., 2005; Cohen et al., 2012; Watabe-Uchida et al., 2017; Berke, 2018; Jeong et al., 2022). This foundational proposal is built on the finding that activity of dopamine neurons at the time a predictive stimulus is encountered is directly relative to the amount and probability of reward predicted by that stimulus (Fiorillo et al., 2003; Tobler et al., 2005). Expected value encoding in dopamine neurons is thought to be incrementally acquired over the course of learning, ultimately reflecting a stable estimate that is used to drive behavior (Fiorillo et al., 2003; Tobler et al., 2005; Hart et al., 2015; Keiflin and Janak, 2015; Mendoza et al., 2019).

Expected value encoding, however, does not well capture the exquisite flexibility of behavioral responses to adapt to changing internal and external states (Robinson and Berridge, 2013; Dayan and Berridge, 2014; Cone et al., 2016). For instance, through alterations in homeostatic processes one can crave intensely salty seawater that is normally aversive and immediately exhibit attraction to cues predictive of receipt of that saltwater on their first encounter, without the need for incremental learning to occur (Richter, 1936; Fudim, 1978; Stouffer and White, 2005; Robinson and Berridge, 2013). If expected value encoding in dopamine neurons is responsible for scaling our behavior to environmental cues, then it is not clear how can one reconcile this elaborate and slow process with instant revaluation of learned cues. However, it has remained difficult to resolve this discrepancy, as approaches to drive such revaluations generally require dramatic changes in physiological state, which can obscure the underlying neurobiological mechanisms (Cone et al., 2016; Fortin and Roitman, 2018; Hsu et al., 2020; Grove et al., 2022). As a result, it remains unresolved whether expected value encoding is a feature of dopamine systems that is critical for adaptive behavior or merely a byproduct of the generally stable, controlled behavior used in prior studies.

To resolve this long-standing issue we used a behavioral phenomenon called occasion setting, where the long-running expected value of a conditioned stimulus is dissociated from its trial-to-trial value as the relation between this stimulus and reward is predicated on the prior presence of a separate, distinct stimulus (Holland, 1992; Fraser and Holland, 2019; Fraser and Janak, 2019). Using a variety of complementary approaches, we reveal that activity within the mesolimbic dopamine system and its projection to the nucleus accumbens is necessary for the incorporation of changes in expected value to drive behavioral adaptation. We find that occasion setters dynamically alter expected value encoding in mesolimbic dopamine release and accumbens neural encoding, on a trial-by-trial basis. These data provide new evidence that, as opposed to generating stable, long-running estimates of value, the mesolimbic dopamine system rapidly updates value in response to changing sensory inputs to generate flexible cue-motivated reward-seeking.

## RESULTS

### Occasion setting reflects rapid value updating and engages model-based control

To dissociate stable value encoding from the rapid revaluation of the motivation attributed to cue, we took advantage of a behavioral process called occasion setting. Occasion setting is a general behavioral phenomenon where the predictive relationship between a conditioned stimulus (CS) and reward is hierarchically gated by a separate and non-overlapping discrete cue, an occasion setter (OS; **Figure 1A**). Occasion setters can be thought of as analogous to contexts but allow for experimenter control over their presentation and actions, for precise assessment of how they can dynamically alter behavior. In our task, the CS’s reward predictive value varies on a trial-by-trial basis, toggling between *P*=1.0 for OS-⟶CS trials and *P*=0.0 for CS alone trials. This process results in behavioral responding being highest to the CS on trials when the OS had preceded its presentation (**Supplemental Figure 1**). We characterized the mechanisms of occasion setters to confirm their actions are to rapidly modulate the value of conditioned stimuli. First, we extinguished responding to the occasion setter to demonstrate that the high responding on trials where the OS preceded the CS is solely driven by the occasion setting modulating the conditioned stimulus, rather than the OS itself predicting reward. Following extinction, there was no longer responding to the occasion setter alone, but the ability of the occasion setter to update the value of the conditioned stimulus remained unaltered (a new analysis of data from Fraser & Janak 2019; **Figure 1B**; effect of trial type F_(2,76)_=101, p<0.0001; effect of extinction F_(1,38)_=28.12, p<0.0001; interaction of trial type and extinction F(2,76)=3.476, p<0.0359; in each group OS⟶CS vs CS alone p<0.01 and OS⟶CS vs. OS alone p<0.0001). In other words, occasion setting is not driven by summation of probabilities nor does occasion setting reflect trace conditioning to the occasion setting cue. We also observed that occasion setters themselves are valuable and also transform conditioned stimuli to become valued and this process, too, is resistant to extinction (**Figure 1C**; effect of trial type F_(2,75)_=9.605, p<0.001; effect of extinction F_(1,38)_=0.1934, p=0.6626; OS⟶CS vs. CS alone p<0.0001; OS alone vs CS alone p=0.0047). Collectively, it is the actions of occasion setters to modulate the value of conditioned stimuli that makes them desirable and drives reward-seeking.

**Figure 1.**
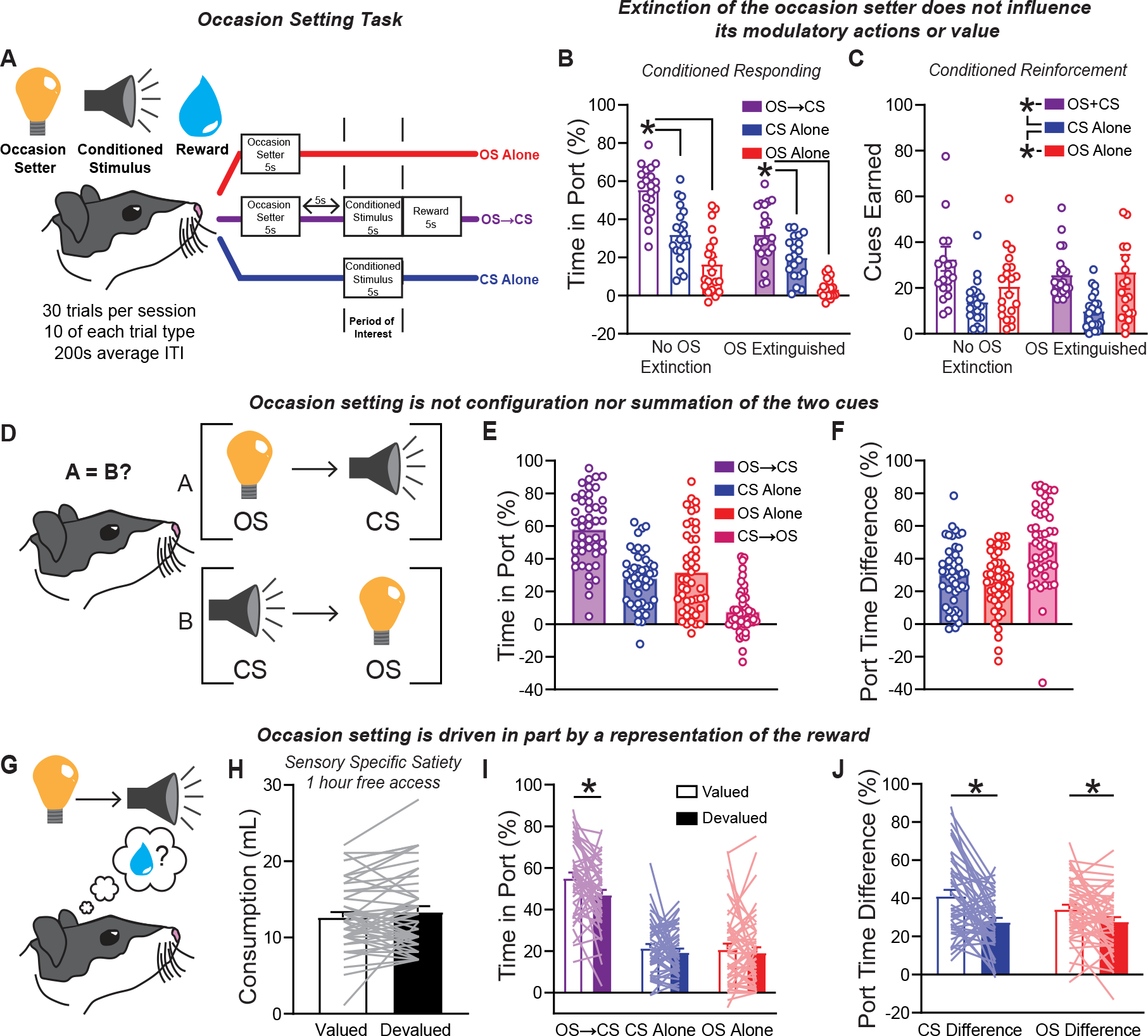
Occasion setting reflects a psychological process that involves the rapid modulation of reward expectation. A. Overview of the occasion setting task. A light served as the occasion setter, white noise as the conditioned stimulus, and 15% sucrose was the reward. Rats were presented with three trial types per session, the occasion setter alone, the conditioned stimulus alone, or the occasion setter followed by a gap then the conditioned stimulus and reward. B. Conditioned responding as assessed by percent time in port before and after occasion setter extinction (*new analysis of data from Fraser & Janak 2019*). C. Conditioned reinforcement (*new analysis of data from Fraser & Janak 2019*). D. Overview of the potential for configuration to explain occasion setting. E. Conditioned responding during the test for configural learning with CS⟶OS probe trials (n=46). F. Difference in responding on OS⟶CS trials relative to CS alone, OS alone, and CS⟶OS trials. G. Schematic of potential for reward-seeking to be driven by a representation of the reward to be earned. H. Reward consumption in mL for the one-hour pre-session access (n=46). I. Conditioned responding following sensory specific satiety. J. Difference in responding on OS⟶CS trials relative to CS alone and OS alone trials following sensory specific satiety. OS, occasion setter. CS, conditioned stimulus. ^*^p<0.05.

We next wanted to confirm that this contextual cue, the occasion setter, updates the value attributed to the conditioned stimulus and that this is not due to configuration or summation of the two stimuli into one “super” stimulus (**Figure 1D**). If rats were configuring, then we would expect that altering the order of presentation of cues would have no effect. It is critical to rule out configuration to confirm that the occasion setter is updating the value attributed to the conditioned stimulus, not contributing to a novel representation. Moreover, given that the average probability of each cue in our task was equal (*P*=0.50), we also wanted to rule out the potential for summation of the probabilities of OS⟶CS trials determining the increased reward-seeking we observed. To test this, we asked whether rats configured the occasion setter and conditioned stimulus, by reversing their order. During this probe test we observed little responding to the occasion setter when it was preceded by the conditioned stimulus on reversed order probe trials (**Figure 1E**; effect of trial type F_(3,45)_=71.44, p<0.0001; OS⟶CS vs. CS⟶OS p<0.0001; **Figure 1F**; effect of discrimination F_(2,45)_=23.72, p<0.0001; CS⟶OS vs. CS discrimination p<0.0001; CS⟶OS vs OS discrimination p<0.0001). The lack of any behavioral response to the occasion setter when it now appeared after the conditioned stimulus supports that rats are not engaging in configuration. Overall, the actions of occasion setters are to dynamically alter the value of a conditioned stimulus on a tri-al-to-trial basis.

Finally, we explored what psychological mechanisms might explain the increased responding on OS⟶CS trials. We wanted to dissociate whether rats simply updated their value estimate, a model-free process, or if in addition they increased responding in part by evoking a representation of the sucrose reward that they would earn on these trials, referred to as a model-based process (Dayan and Berridge, 2014; **Figure 1G**). To test these two possibilities, we made use of sensory specific satiety where the rats had free access the reward they would earn, 15% sucrose (devalued condition), or of equal reward, 15% maltodextrin (valued condition), for one hour. In these free access sessions, there was no difference in the amount of reward consumed for either valued or devalued conditions (**Figure 1H**; paired t-test t_45_=1.383, p=0.1735). Immediately following free access consumption rats were placed in behavioral chambers and tested under extinction conditions (no reward delivered) to see if there was a selective effect of devaluing the reward typically earned. We found that devaluation resulted in a selective deficit in responding only on OS⟶CS trials (**Figure 1I**; interaction of devaluation and trial type F_(2,90)_=3.382, p=0.0383; OS⟶CS p=0.0003; CS alone p=0.6739; OS alone p=0.8223). Consequently, devaluation resulted in a reduction in the ability of rats to discriminate when they should expect reward following the conditioned stimulus (**Figure 1J**; effect of devaluation F_(1,45)_=13.63, p=0.0006; interaction of discrimination and devaluation F_(1,54)_=4.212, p=0.046; CS difference p<0.0001; OS difference 0.0217). This supports the notion that the occasion setter acts in part by generating an expectation of the precise characteristics of the reward at the time of the conditioned stimulus in addition to altering the value of this cue.

### Phasic VTA dopamine neuron activity is necessary for contextual modulation of motivation

The ability to make flexible inferences about cues has been shown to be mediated in part by dopamine neurons residing in the ventral tegmental area (Chang et al., 2017; Sharpe et al., 2017, 2020; Keiflin et al., 2019; Maes et al., 2020; Kutlu et al., 2022). However, evidence for these contributions come from behavioral tests in which the value attributed to cues is stable, which can be accounted for by traditional models of reinforcement learning that produce value estimates that are slow to acquire and resistant to change. We wondered if dopamine neurons would be essential for mediating cue-driven behavior if we selectively eliminated their activity during a contextual cue that would shift the state of what is valuable in the world. To do so, we trained male and female TH-cre rats with either the inhibitory green-light activated opsin halorhodopsin (Halo) or a control GFP virus selectively expressed in dopamine neurons (**Figure 2A-C**; n=12 per group; 6 males and 6 females per group) as above. There were no differences in the behavior of TH-cre rats with Halo or GFP expressed in the ventral tegmental area (VTA) during training (**Supplemental Figure 1**). To assess the functions of VTA dopamine neurons we delivered 5.5s of 532-nm light bilaterally into the VTA during discrete task epochs, taking advantage of the ability to separate out contextual processing from cue processing in our task. We first demonstrated that inhibition of VTA dopamine neuron outside of task events had no effect on behavior (**Figure 2D**). As expected, both rats with GFP and Halo exhibited higher reward-seeking on OS⟶CS trials than CS alone (**Figure 2E**; effect of trial type F_(1,34)_=55.68, p<0.001; effect of group F_(1,22)_=0.06043, p=0.8081; Halo p=0.0004; GFP p=0.0002) and OS alone trials (Halo p<0.0001; GFP p<0.0001), and the average discrimination for each group of rats in discerning when the CS would be rewarded was similar between groups (**Figure 2F**; effect of group F_(1,22)_=3.760, p=0.5063; effect of difference F_(1,22)_=3.299, p=0.0830).

**Figure 2.**
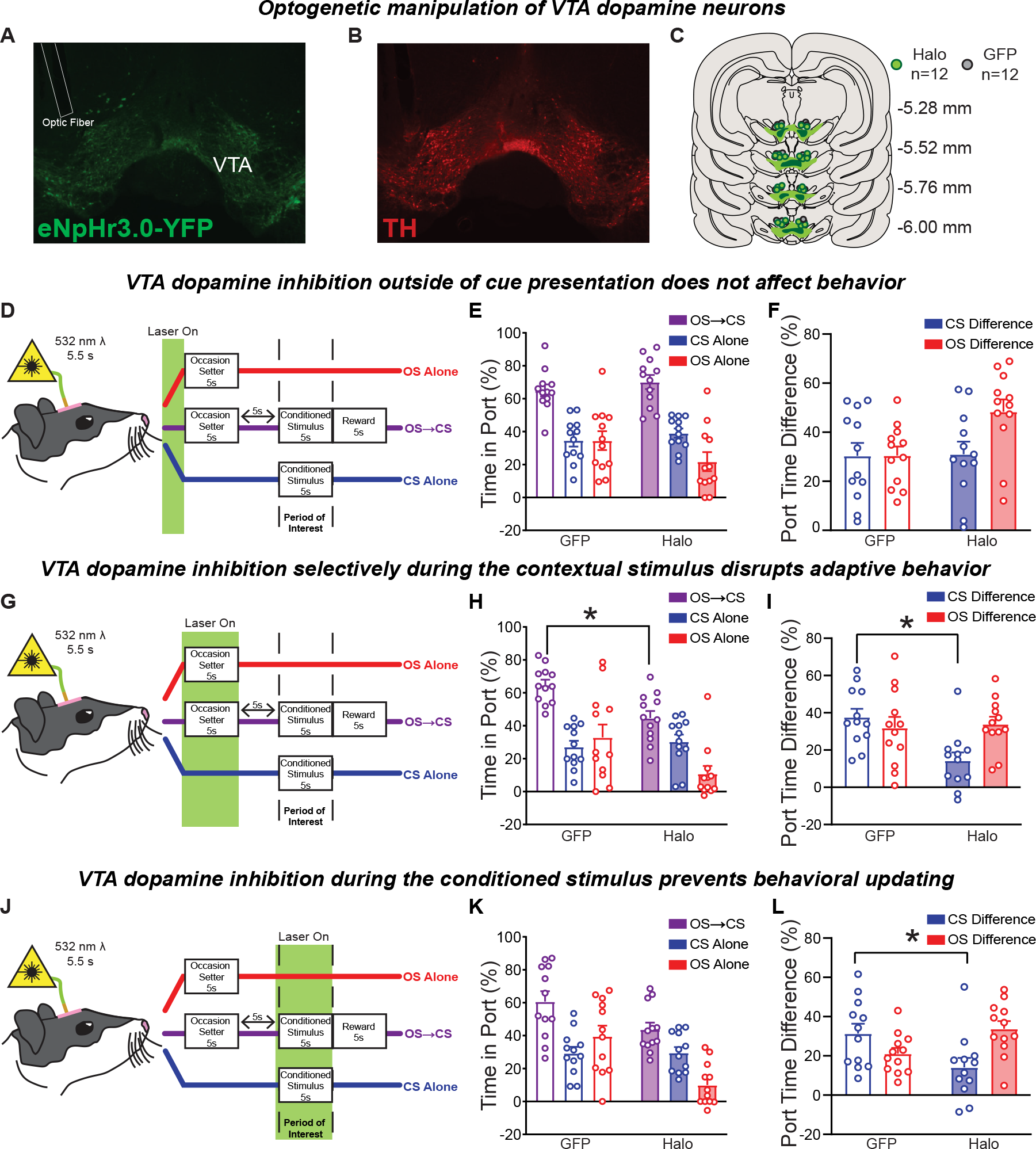
VTA dopamine neurons are necessary for updating the value of reward-paired cues to guide behavior. A. Representative expression of cre-dependent halorhodopsin in the VTA of TH-Cre rats. B. Co-staining of TH for the same image in A. C. Expression mapping of halorhodopsin and the placement of optic fibers for all rats included (n=12 per group). Light green indicates largest area of expression, dark green represents smallest area of expression observed. D. Schematic of pre-trial dopamine neuron inhibition. E. Conditioned responding during occasion setting following 532 nm light delivery into the VTA for 5.5s, 10s prior to every trial. F. Difference in responding on OS⟶CS trials relative to CS alone and OS alone trials following pretrial light delivery. G-I. Same as D-F but when light was delivered for 5.5s beginning 0.5s before the occasion setter. J-L. Same as D-F but when light was delivered for 5.5s beginning 0.5s before the conditioned stimulus. OS, occasion setter. CS, conditioned stimulus. eNpHr3.0, halorhodopsin. TH, tyrosine hydroxylase. ^*^p<0.05.

When we selectively inhibited dopamine neurons during the occasion setter – the context-like cue – rats were no longer able to estimate whether a conditioned stimulus was or was not predictive of reward. VTA dopamine neuron inhibition during the OS period selectively affected reward-seeking for rats expressing Halo (**Figure 2H**; effect of group F_(1,22)_=7.1, p=0.0142; effect of trial type F_(2,44)_=33.02, p<0.0001; interaction F_(2,44)_=5.457, p=0.0076), reducing their reward-seeking on OS⟶CS trials (p=0.0151) and OS Alone trials (p=0.0065) relative to GFP controls. In addition, rats with Halo no longer exhibited a significant elevation in reward-seeking on OS⟶CS trials relative to CS alone trials (p=0.0651). This reduction in reward-seeking resulted in a significant reduction in the ability of rats with Halo to discriminate between trials in which the CS was reinforced versus when it was not reinforced, relative to GFP controls (**Figure 2I**; interaction F_(1,22)_=5.080, p=0.0345; CS difference p=0.0025; OS difference p=0.9465).

Finally, we assessed whether VTA dopamine neuron activity during the conditioned stimulus – the cue that evokes motivated reward-seeking – was necessary. When VTA dopamine neurons were inhibited during the CS, reward-seeking for Halo rats was impaired (**Figure 2K**; effect of group F_(1,22)_=7.375, p=0.0126; effect of trial type F_(1,37)_=40.54, p<0.0001; interaction F_(2,44)_=10.58, p=0.0002). However, rats still exhibited the expected behavioral pattern (within-group OS⟶CS responding was higher than CS alone and OS alone trials; post hoc p’s<0.05) with no differences between groups. Despite this, VTA dopamine neuron inhibition significantly impaired the ability of Halo rats to properly estimate when the CS would be rewarded relative to GFP rats (**Figure 2L**; interaction F_(1,22)_=16.25, p=0.0006; CS difference p=0.0129; OS difference p=0.0813). There were no effects of VTA dopamine neuron inhibition during reward delivery, the trace between the OS and CS, or if inhibition was not constant during the OS (**Supplemental Figure 2**).

Taken together, VTA dopamine neurons are necessary for the processing of contexts and the future utilization of contextual information to rapidly update the value of an ambiguous cue. This provides strong evidence that the function of VTA dopamine neurons is to rapidly compute the immediate motivational value of reward-paired cues to drive the appropriate degree of reward-seeking.

### Nucleus accumbens neural activity and dopamine release are necessary for the utilization of updated value to guide reward-seeking

VTA dopamine neurons project strongly to the nucleus accumbens, where their actions have been implicated in encoding the expected value of cues to guide behavior (Day et al., 2007; Flagel et al., 2011; Clark et al., 2013; Hart et al., 2015; Keiflin and Janak, 2015; Aitken et al., 2016; Saunders et al., 2018). We next investigated the neurochemical mechanisms in the nucleus accumbens that underlie occasion setting using intracranial neuropharmacology (**Figure 3A**). Flupenthixol, a general dopamine receptor antagonist, delivered into the nucleus accumbens core resulted in an expected small but significant decreases in intertrial port entries (Figure 3D; t_11_=4.874, p=0.0005) and intertrial port time (**Figure 3E**; t_11_=4.705, p=0.0006). We found that blocking dopamine signaling significantly reduced context-modulated reward-seeking (**Figure 3B**; effect of trial type F(1,21)=29.58, p<0.0001; effect of drug F(1,11)=51.65, p<0.0001; interaction F(1,20)=4.952, p=0.0198). Dopamine antagonism reduced reward-seeking on all trials relative to saline (OS+CS p=0.006; CS alone p=0.0028; OS alone p=0.0008) and resulted in rats being unable to elevate reward seeking on OS⟶CS trials relative to CS alone trials (p=0.3447). However, even with potentially reducing overall locomotor activity, nucleus accumbens core dopamine antagonism significantly reduced overall discrimination between reinforced and non-reinforced trials in the task (**Figure 3C**; effect of drug F_(1,11)_=12.36, p=0.0048; within each discrimination p<0.05). Thus, not only is the activity of VTA dopamine neurons essential for occasion setting, but so is dopamine release in the nucleus accumbens.

**Figure 3.**
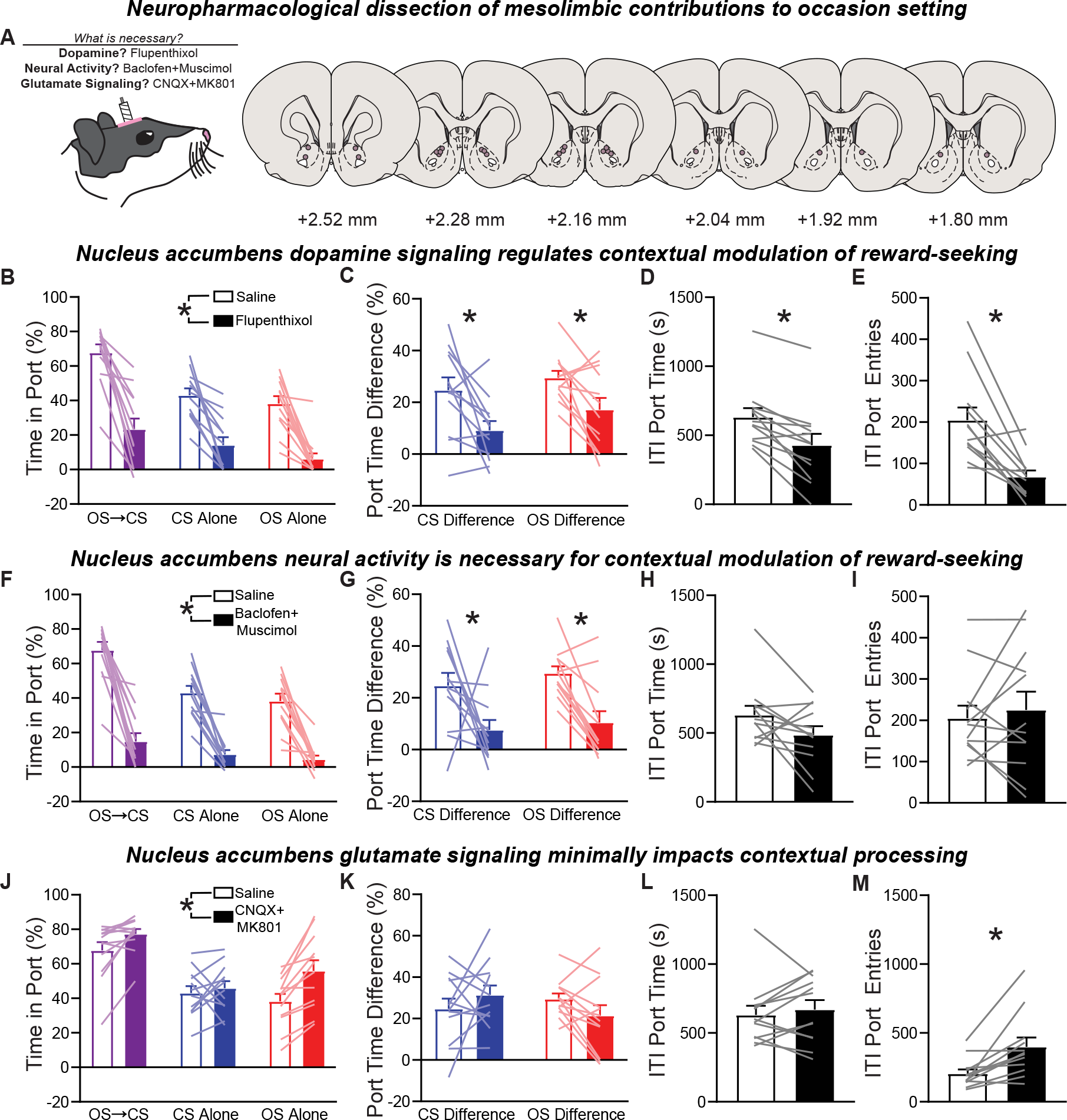
Dopamine release and neural activity within the nucleus accumbens are required for the contextual modulation of value to guide reward-seeking. A. Schematic of general approach and recreation of microinjector tips within the nucleus accumbens core (n=12). B. Conditioned responding as assessed by percent time in port following saline and flupenthixol infusions in the nucleus accumbens core. C. Difference in responding on OS⟶CS trials relative to CS alone and OS alone trials following flupenthixol infusion. D. Intertrial port time following flupenthixol. E. Intertrial port entries following flupenthixol. F-I. Same as B-E but following infusion of baclofen and muscimol into the nucleus accumbens core. J-M. Same as B-E but following infusion of CNQX and MK801 into the nucleus accumbens core. OS, occasion setter. CS, conditioned stimulus. ^*^p<0.05.

Nucleus accumbens neural activity is linked to the encoding of Pavlovian reward-paired cues and is shaped by dopamine release (Day et al., 2006; Ambroggi et al., 2011; du Hoffmann and Nicola, 2014). We focally inactivated the nucleus accumbens core via infusion of the GABA agonists baclofen and muscimol. Transient inactivation of the nucleus accumbens core reduced reward-seeking compared to behavior under saline (**Figure 3F**; effect of trial type F_(1,18)_=33.78, p<0.0001; effect of drug F_(1,11)_=92.15, p<0.0001; interaction F_(1,13)_=6.108, p=0.0220). Inactivation significantly reduced time in the port on OS⟶CS (p=0.0001), CS alone (p=0.0001), and OS alone trials (p=0.003) relative to saline. In contrast to significantly higher port time on OS+CS trials relative to CS alone (p=0.0058) and OS alone trials (p<0.0001) following saline, there was no difference in time on OS-⟶CS trials relative to CS alone (p=0.7028) or OS alone trials (p=0.4240) following inactivation. As a result, inactivation reduced the ability to rats to discriminate amongst reinforced trials and CS alone and OS alone trials (**Figure 3G**; effect of drug F_(1,11)_=9.974, p=0.0091; within each discrimination p<0.01). Inactivation did not impact intertrial port time (**Figure 3H**; t_11_=1.917, p=0.0816) or intertrial port entries (**Figure 3I**; t_11_=0.5779, p=0.5750), indicating that nucleus accumbens inactivation had a selective impact on cue-evoked responding.

We have previously demonstrated that frontal cortical and basolateral amygdala activity is necessary for occasion setting, both of which project into the nucleus accumbens, and the activity of glutamatergic amygdala inputs to the nucleus accumbens shapes behavioral responses to discriminative cues (Ambroggi et al., 2008; Fraser and Janak, 2023). Thus, we assessed whether functional glutamate signaling in the nucleus accumbens was essential for occasion setting via infusion of the AMPA and NMDA receptor antagonists CNQX and MK801. In the nucleus accumbens core, blockade of

AMPA and NMDA receptors increased overall responding in the occasion setting task (**Figure 3J**; effect of trial type F_(1,19)_=32.47, p<0.0001; effect of drug F_(1,11)_=33.16, p=0.0001; interaction F_(1,17)_=2.904, p=0.0904). Analysis of the impact of drug infusion revealed that glutamate antagonism selectively increased reward-seeking on OS alone trials relative to saline (p=0.0069). This resulted in behavior following glutamate receptor antagonism on OS alone trials being statistically indistinguishable from behavior on OS⟶CS trials following saline infusion (p=0.3524). Despite this, rats still discriminated at a similar level under glutamate antagonism (**Figure 3K**; effect of drug F_(1,11)_=0.0305, p=0.8644) and while there was no impact of treatment on intertrial port time (**Figure 3L**; t_11_=0.6838, p=0.5082) there was a significant elevation in intertrial port entries (**Figure 3M**; t_11_=3.579, p=0.0043). Collectively it appears that glutamate release in the accumbens may constrain aspects of contextual processing, but there is an essential contribution of striatal dopamine and neural activity for the contextual revaluation of expected value driven by occasion setters.

### Nucleus accumbens dopamine release dynamically encodes the motivational value of conditioned stimuli

We have shown that dopamine’s actions in the nucleus accumbens are necessary for the control of reward-seeking by contexts, namely occasion setters, and that VTA dopamine neurons are essential at the time of occasion setters and their conditioned stimuli to support reward-seeking. However, while this evidence is suggestive of a rapid modulation of value encoding on a trial-to-trial basis (e.g., toggling between *P*=1.0 and *P*=0.0 for the CS), it remains possible that dopamine release instead tracks the overall expected value (*P*=0.50) of each of the two cues in this task. To tease apart these potential functions we measured dopamine signaling in the nucleus accumbens core as rats performed the occasion setting task, using in vivo fiber photometry recordings of the genetically-engineered fluorescent sensor dLight1.3b (**Figure 4A-C**; Patriarchi et al., 2018). We analyzed the peak dopamine response following OS and CS presentation as well as after reward delivery. We found that dopamine signaling was robustly elicited by presentation of the context (**Figure 4D-F**) - the occasion setter - which aligned well with our finding that inhibition of dopamine neurons at this timepoint significantly disrupted behavior. Dopamine release was equivalent to the OS regardless of whether the OS was followed by the CS (**Figure 4I**; effect of trial type Q_3_=11.14, p=0.0012; OS⟶CS vs. OS alone p>0.9999). However, dopamine release to the conditioned stimulus was much greater on OS⟶CS trials, where the occasion setter informed this cue would be rewarded, than on CS alone trials (**Figure 4G-H**). Critically, we only observed significantly greater dopamine release in response to the conditioned stimulus on OS⟶CS trials, but not on CS alone trials (**Figure 4J**; effect of trial type F_(1,7)_=143.9, p<0.0001; OS⟶CS vs. CS alone p=0.004; OS⟶CS vs. OS alone p=0.0137; CS alone vs. OS alone p=0.5221). This pattern of results was similar when we analyzed the area under the curve for the total cue length, with no difference for the OS (Figure 4K; effect of trial type Q_3_=11.14, p=0.0012; OS⟶CS vs. OS alone p>0.9999) and a significant difference in dopamine release at the time of the conditioned stimulus on OS⟶CS trials (**Figure 4L**; effect of trial type F_(1,7)_=21.12, p=0.0016; OS⟶CS vs. CS alone p=0.0104; OS⟶CS vs. OS alone p=0.0222; CS alone vs. OS alone p=0.8613). This all-or-nothing encoding of the conditioned stimulus is counter to the expectation that dopamine release tracks the long-running expected value of cues. Instead, this is consistent with a framework that dopamine neuron activity and dopamine release are critical for encoding the current motivational value of stimuli to dynamically scale behavior.

**Figure 4.**
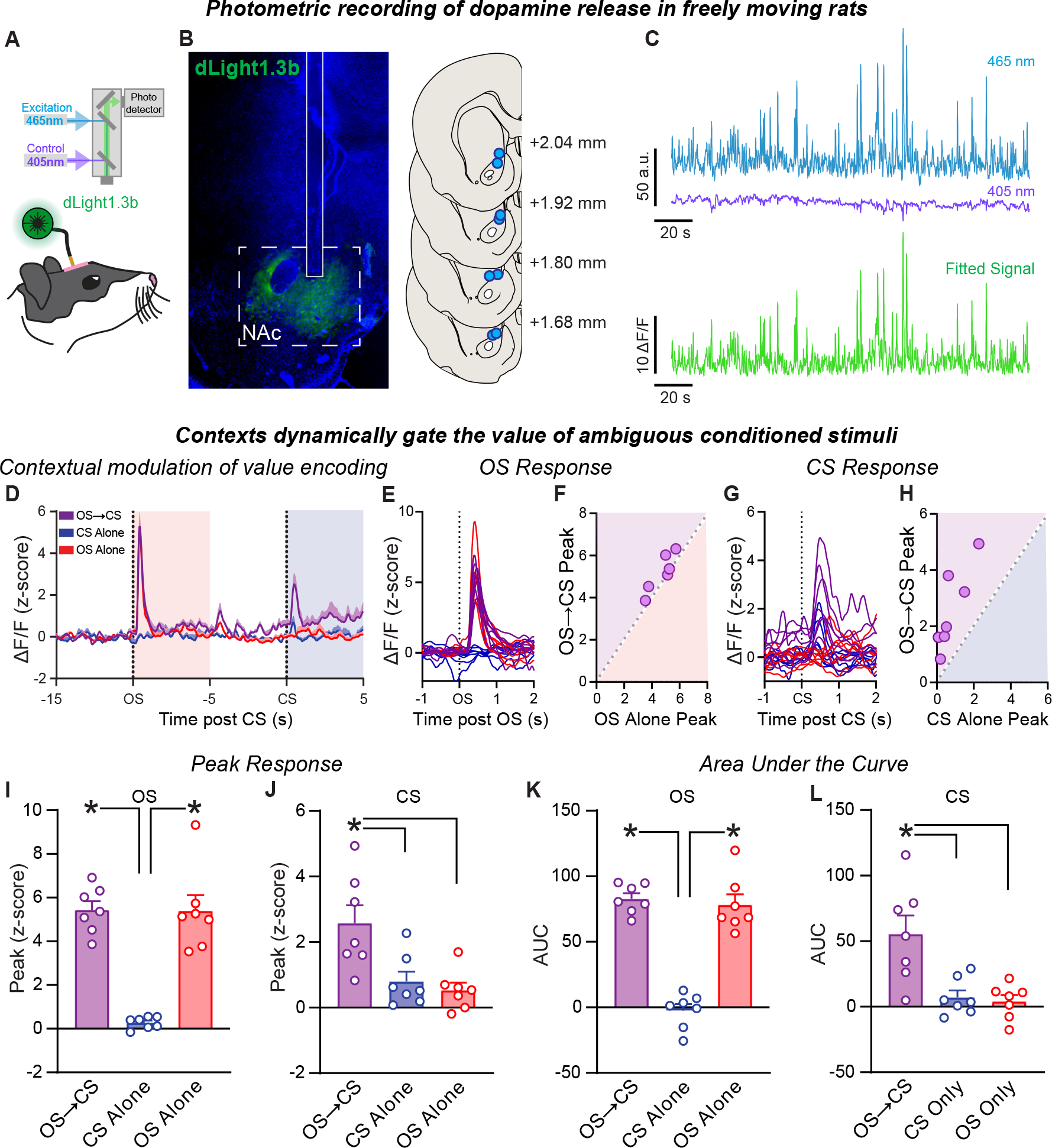
Dopamine release dynamically encodes the current value of reward-associated cues. A. Schematic of recording approach. B. Representative dLight1.3b expression in the nucleus accumbens and recreation of optic fiber placements for all rats included (n=8). C. Representative raw fluorescence emitted by dLight1.3b following 465 nm excitation, isosbestic 405 nm excitation, and resultant ΔF/F produced by fitting and normalizing 465nm fluorescence to 405 nm fluorescence. D. Average dopamine release during occasion setting (n=7). E. Dopamine release following occasion setter presentation across all trials for each rat. F. Peak dopamine release observed for each rat during the occasion setter contrasted between OS⟶CS and OS alone trials. G. Dopamine release following conditioned stimulus presentation across all trials for each rat. H. Peak dopamine release for each rat during the conditioned stimulus contrasted between OS⟶CS and CS alone trials. I. Peak dopamine release (ΔF/F) during the occasion setter, or the period when it would be presented, across all trial types. J. Peak dopamine release (ΔF/F) during the conditioned stimulus, or the period when it would be presented, across all trial types. K-L. Same as I-J but for the quantification of area under the curve during each interval. OS, occasion setter. CS, conditioned stimulus. ^*^p<0.05.

We observed that dopamine release to the contextual cue, the occasion setter, was greater than that to the CS. This could be a reflection of back-propagation of value from the most proximal to most distal predictors of reward availability, consistent with reward-prediction error theories of dopamine function (Schultz et al., 1997; Hollerman and Schultz, 1998; Collins et al., 2016; Sutton and Barto, 2018; Mohebi et al., 2019; Amo et al., 2022). To tease this apart, we explicitly manipulated the potential value attributed to the occasion setter via extinction (e.g., presenting only the occasion setter repeatedly without any other stimuli or reward; Fraser and Janak, 2019). This manipulation has no effect on the discrimination achieved by rats (for example see **Figure 1B**). In turn, we saw no general effect of this manipulation on dopamine release (**Figure 5A-E**). There was no effect of OS extinction on the response to the occasion setter (Figure 5F, peak; effect of extinction F_(1,7)_=0.1204, p=0.7388; interaction of extinction and trial type F_(1,7)_=0.1486, p=0.8636; **Figure 5H**, AUC; effect of extinction F_(1,7)_=1.468, p=0.2649; interaction of extinction and trial type F_(1,7)_=0.6199, p=0.5558) nor to the conditioned stimulus (**Figure 5G**, peak; effect of extinction F_(1,7)_=0.9495, p=0.3623; interaction of extinction and trial type F_(1,7)_=0.1740, p=0.8426; **Figure 5I**, AUC; effect of extinction F_(1,7)_=2.049, p=0.1954; interaction of extinction and trial type F_(1,7)_=0.3803, p=0.6923). Consequently, it does not appear that trace conditioning or backpropagation of value to the occasion setter is directly responsible for its ability to gate the value of a conditioned stimulus.

**Figure 5.**
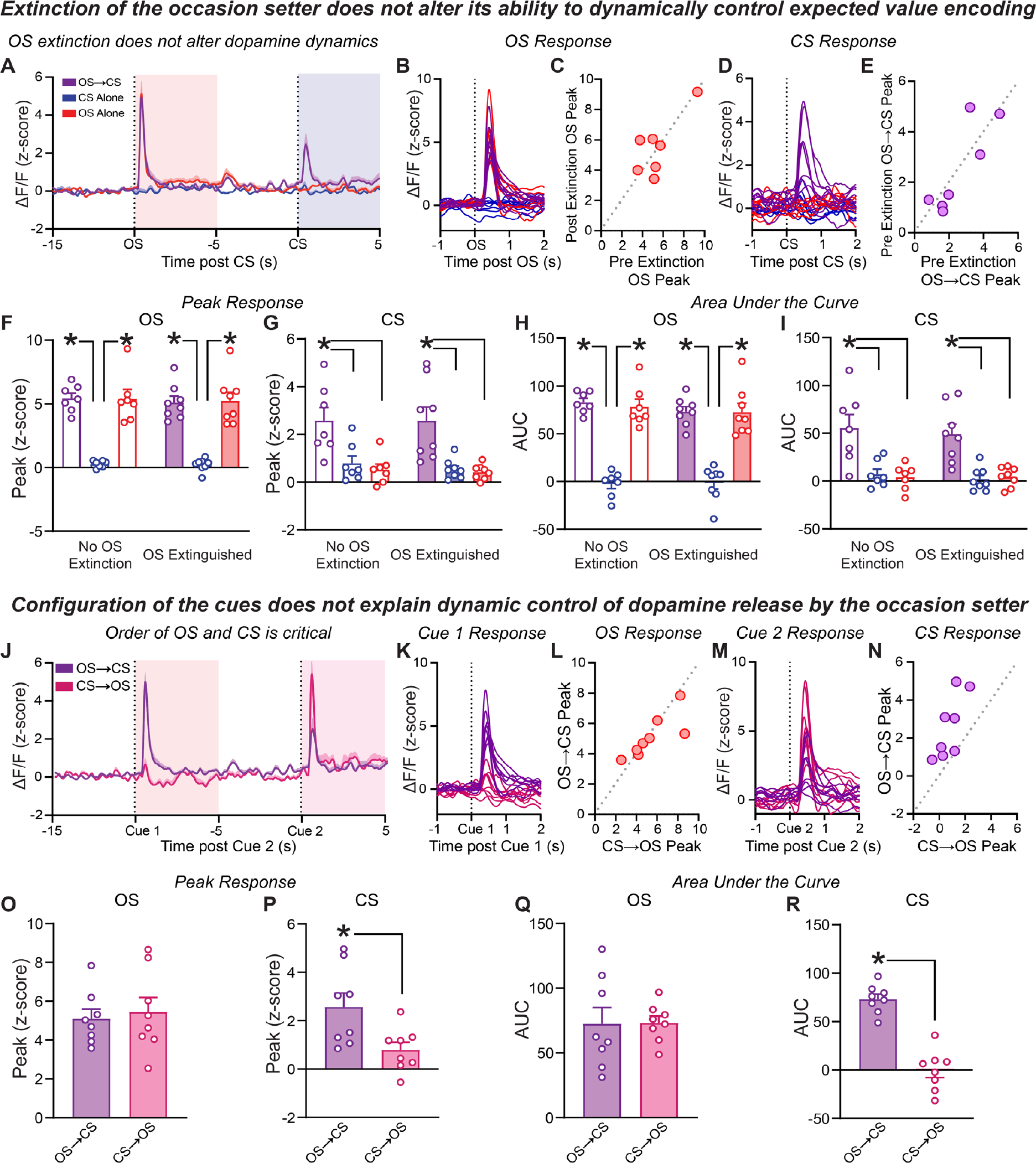
Dopamine release in the nucleus accumbens reflects the ability of occasion setters to dynamically regulate expected value. A. Average dopamine release during occasion setting after occasion setter extinction (n=8). B. Dopamine release following occasion setter presentation after occasion setter extinction across all trials for each rat. C. Peak dopamine release observed for each rat during the occasion setter contrasted for OS alone trials before and after occasion setter extinction. D. Dopamine release following conditioned stimulus presentation across all trials for each rat following occasion setter extinction. E. Peak dopamine release for each rat during the conditioned stimulus contrasted for OS⟶CS trials before and after occasion setter extinction. F. Peak dopamine release during the occasion setter for all three trials before and after occasion setter extinction. G. Peak dopamine release during the conditioned stimulus for all three trials before and after occasion setter extinction. H. Average dopamine release during the occasion setter as assessed by the area under the curve before and after occasion setter extinction. I. Average dopamine release during the conditioned stimulus as assessed by the area under the curve before and after occasion setter extinction. J. Average dopamine release during occasion setting on OS⟶CS trials relative to configural CS⟶OS probe trials (n=8). K. Dopamine release to the first cue presented, the OS for OS⟶CS trials or the CS for CS⟶OS probe trials. L. Peak dopamine release observed for each rat to the occasion setter presented either in sequence (OS⟶CS) or out of sequence. (CS⟶OS). M. Dopamine release to the second cue presented, the CS for OS⟶CS trials or the OS for CS⟶OS probe trials. N. Peak dopamine release for each rat during to the conditioned stimulus contrasted presented either in sequence (OS⟶CS) or out of sequence (CS⟶OS). O. Peak dopamine release during the occasion setter (cue 1 for OS⟶CS and cue 2 for CS⟶OS). P. Peak dopamine release during the conditioned stimulus (cue 2 for OS⟶CS and cue 1 for CS⟶OS). Q-R. Same as O-P but average dopamine release as assessed by area under the curve. OS, occasion setter. CS, conditioned stimulus. ^*^p<0.05.

We wanted to confirm that rapid modulation of dopamine encoding of value is due to contextual control by the occasion setter. To do so we tested in the same rats whether altering the order of the cues could explain the observed pattern of dopamine release (but see the lack of impact of this manipulation on behavior as in **Figure 1H-J**). Given each cue in this task has the same probability of predicting reward (*P*=0.50) it was critical to rule out whether the mere ordering of any two stimuli on a trial would drive a difference in dopaminergic encoding of the second cue presented. However, when we tested this by presenting the CS before the OS, there was no modulation of dopamine to the OS as compared to on standard OS⟶CS trials, supporting the notion that summation of probability does not explain occasion setting (**Figure 5J-N**). There was no difference in the peak dopamine response (**Figure 5O**; t_7_=0.7579, p=0.4732) to the occasion setter whether it preceded the conditioned stimulus (OS⟶CS trials) or when it occurred after the conditioned stimulus (CS⟶OS trials). While the occasion setter dynamically gated dopamine release to the conditioned stimulus on OS⟶CS trials as expected, the degree of dopamine release to the conditioned stimulus was akin to that observed on CS alone trials (**Figure 5P**; CS⟶OS vs OS⟶CS t_7_=4.511, p=0.0028; CS⟶OS vs CS alone CS t_7_=0.9408, p=0.3781). There was no effect on the amount of dopamine release to the occasion setter either as assessed by area under the curve (**Figure 5Q**; t_7_=0.0807, p=0.9379). Dopamine release to the conditioned stimulus was overall greater on OS⟶CS trials than on CS⟶OS trials (**Figure 5R**; t_7_=7.24, p=0.0002). Together, these data show nucleus accumbens dopamine signals encode contextual, or higher-order, information that is rapidly utilized to dynamically alter the value attributed to reward-paired cues and scale reward-seeking appropriately. This gating of expected value is consistent with the integration of contextual information to rapidly disambiguate the immediate motivational value of reward-paired cues to in turn drive appropriate reward pursuit.

### Contexts dynamically alter nucleus accumbens neural encoding of reward-paired cues

Finally, we wanted to understand whether the alterations observed in dopamine encoding of value are reflected in the dynamics of downstream striatal neural activity. Reversible inactivation of the nucleus accumbens significantly disrupted behavior in this task, suggesting that striatal encoding of these stimuli is essential for the contextual modulation of behavior (**Figure 3F-I & Supplemental Figure 3**). To assess this, we implanted rats with microdrives of 16 tungsten wires in the nucleus accumbens core and recorded neural activity (n=235 neurons) as they performed the occasion setting task (**Figure 6A-B**). We were most interested in understanding if neural encoding of the conditioned stimulus was altered by the context in which it was encountered, controlled by the separate, prior presentation of the occasion setter. To dissect neural encoding of the task, we implemented a kernel regression approach to detect significant modulation of individual neurons’ activity by the OS and CS, as well as selective modulation of the CS response by OS presentation. We observed that 69% of recorded nucleus accumbens neurons were significantly modulated by OS presentation (OS-responsive) and 32% were significantly modulated following CS presentation, irrespective of OS presentation (CS-general). On the other hand, 22% of nucleus accumbens neurons demonstrated CS modulation that discriminated based on context (CS-context modulated; **Figure 6C**). Plotting the activity of modulated neurons revealed a mixture of cue-evoked excitations and inhibitions among OS-responsive and CS-general neurons that were indeed stable across trial type and context-independent (**Figure 6D-E,G-J)**.

**Figure 6.**
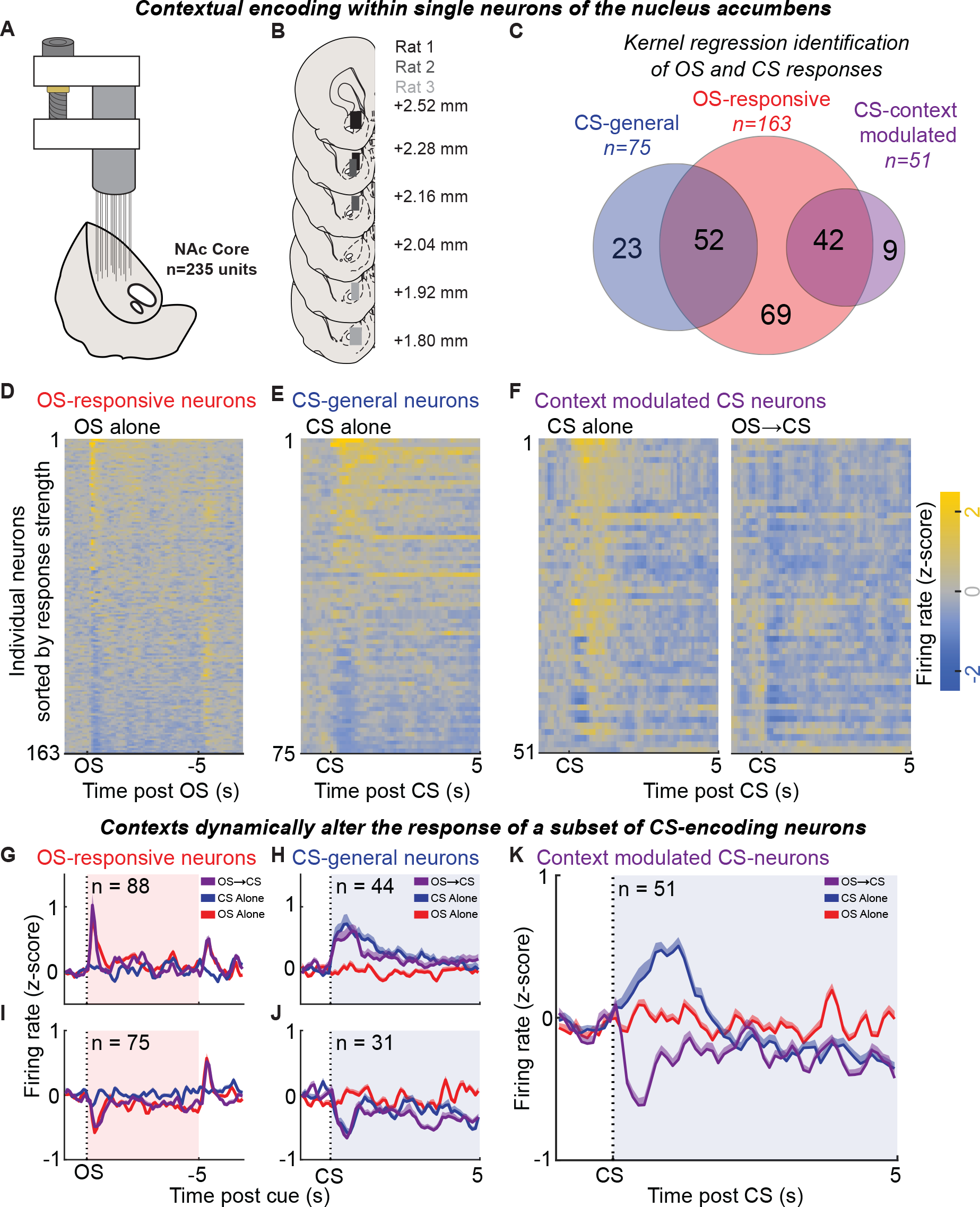
Contextual control of nucleus accumbens encoding of value. A. Schematic of microdrive-based recording approach. B. Recreation of the extent of recording sites within the nucleus accumbens (n=3; 1 male, 2 females). C. Venn diagram of neurons with significant responses to the OS or CS, with distinction for responses to all CS trials (CS-general) or discriminating according to occasion setter (CS-context modulated). D. Heat map of z-score firing rates for OS-responsive neurons on OS alone trials, sorted by their response to OS onset. E. Heat map of z-score firing rates for CS-general neurons on CS alone trials, sorted by their response to CS onset. F. Heat map of z-score firing rates for CS-context modulated neurons on CS alone (left)(left) and OS⟶CS trials (right), sorted by the difference in activity between these two conditions. G. Average (+SEM) response of neurons that were significantly excited by OS onset. H. Same as G, for CS-general neurons excited by CS onset. I-J. Same as G-H for neurons inhibited by cue onset. K. Average (+SEM) response of all context modulated CS-neurons. OS, occasion setter. CS, conditioned stimulus.

We next investigated the response patterns of the CS-context modulated nucleus accumbens neurons. These neurons demonstrated clear differences in firing rate on trials where the CS was presented alone versus trials where the CS was preceded by the OS (**Figure 6F**). Interestingly, the vast majority (49/51) had greater firing rate for CS alone trials, when this cue did not predict reward, than for the CS following OS presentation, when this cue predicted reward. The average activity of these CS-context modulated neurons deflected above baseline for CS alone trials and below baseline for OS⟶CS trials (**Figure 6K**), demonstrating bidirectional contextual control of nucleus accumbens firing patterns. As a result, not only do nucleus accumbens neurons exhibit contextual modulation of cue-encoding, but a context can flip the response sign of accumbens neurons from excitation to inhibition in accordance with the immediate expected value of that cue. These results indicate that rapid revaluation of the value of reward-paired cues comprises alterations in not only dopaminergic encoding of these stimuli, but also in the downstream striatal network.

## DISCUSSION

Dopamine neurons in the midbrain are well-known for their ability to signal prediction errors – the difference between cue-elicited expectations and the outcome that was received (Rescorla and Wagner, 1972; Schultz and Dickinson, 2000; Keiflin and Janak, 2015; Watabe-Uchida et al., 2017; Gardner et al., 2018). Importantly, the degree of cue-elicited excitation in midbrain dopamine neurons is sensitive to the probability and magnitude of reward predicted by a given stimulus, leading to the notion that cue-elicited responses in VTA dopamine neurons reflect the expected value of a given reward-associated stimulus (Fiorillo et al., 2003; Tobler et al., 2005; Cohen et al., 2012). Despite this, a link between expected value and cue-elicited behavior has remained unclear, as freely-behaving animals can exhibit similar levels of conditioned approach for cues of varying expected values (Clark et al., 2012; Dayan and Berridge, 2014; Hart et al., 2015; Mendoza et al., 2019). Exploiting occasion setting, we were able to dissociate long-running expected value from the immediate motivational value of a conditioned stimulus and observe a behavioral report that rats had an accurate estimation of the likelihood of reward receipt. Interestingly, we find that in this situation, where the current value of a conditioned stimulus is dynamically shaped, mesolimbic dopamine release encoded the value of that cue only when it was predictive of reward, and not its average value. Accordingly, inhibition of VTA dopamine neurons selectively during the informative, contextual stimulus was able to prevent the rapid revaluation of cue value. Moreover, dopamine signaling and signal neuron striatal activity were dynamically altered based on immediate value, not the expected value, of the conditioned stimulus. These findings suggest the contribution of phasic VTA dopamine neuron activity to behavior is not necessarily signaling the long-running expected value of cues, but in utilizing a much more flexible and rapidly generated value signal to guide cue-triggered reward-seeking.

We provide evidence that occasion setters act in part by transforming the representation evoked by a conditioned stimulus to one that contains specific information about the reward to be earned, a model-based computation (Dayan and Berridge, 2014). Occasion setting is not driven by summation of expected value, nor does it involve trace conditioning nor configuration of two stimuli (Holland, 1992; Fraser and Holland, 2019). In turn, we only observed dopamine release to a reward-predictive cue when contextual information indicated that this cue was desirable. This contrasts with theories of dopamine function that ascribe dopamine release as encoding the expected value of the conditioned stimuli (Schultz and Dickinson, 2000; Fiorillo et al., 2003; Tobler et al., 2005; Watabe-Uchida et al., 2017; Gardner et al., 2018). How then can we reconcile the ability of occasion setters to rapidly alter value with the long-standing evidence supporting a role for dopamine in expected value encoding? One suggestion is that occasion setters act as a scaling factor in the temporal difference prediction error encoded by dopamine neurons (Zhang et al., 2009). In fact, it has been well-demonstrated that dopamine release in the accumbens can be influenced by alterations in homeostasis, as dopamine release to water-paired cues increases if animals are thirsty or increases in release to food-cues if hungry (Cone et al., 2016; Fortin and Roitman, 2018; Hsu et al., 2020; Grove et al., 2022). However, these state-level manipulations can result in alterations in dopamine activity due to their direct effect on hypothalamic and homeostatic circuits that can more generally inflate the value of cues in the environment (Balleine, 1994; Corbit et al., 2007; Grove et al., 2022). The scaling of dopamine signals in the nucleus accumbens suggests that a function of dopamine neurons is to dynamically encode the immediate behavioral relevance of reward-paired cues (Berridge, 2007, 2012; Smith et al., 2011; Robinson and Berridge, 2013; Dayan and Berridge, 2014; Aitken et al., 2016). The selective gating effect of contexts on the motivational value of cues, and in turn dopamine release and striatal network activity, suggests a role for the mesolimbic dopamine system in encoding and utilizing information about the causal structure of the world to selectively direct behavioral responding to the appropriate target (Chang et al., 2017; Sharpe et al., 2017; Starkweather et al., 2017; Langdon et al., 2018; Gershman and Uchida, 2019; Keiflin et al., 2019).

Occasion setting is a process that captures the control of behavioral responding by contexts (Grahame et al., 1990; Holland, 1992; Holland and Bouton, 1999; Bouton et al., 2006; Fraser and Holland, 2019; Valyear et al., 2023). Despite evidence that dopamine neurons and their striatal targets are essential for the contextual-control of reward-seeking it has been difficult to disentangle the contribution of dopamine neuron activity to encoding contextual control or merely the generation of reward-seeking by cues. This is because traditional context-based approaches (e.g. renewal and reinstatement) are not amenable to separating these processes because the use of a large, multidimensional, and temporally diffuse setting prevents experimenter control over its actions (Crombag et al., 2008; Chaudhri et al., 2010; Floresco, 2015; Valyear et al., 2020, 2023). By reducing a “context” to a brief and discrete event, an occasion setter, we were able to resolve that dopamine neuron activity during the presentation of this context-like occasion setting cue was necessary for that cue to then gate reward-seeking to an ambiguous conditioned stimulus. The necessity of dopamine neuron activity and dopamine release at the time of these contexts, and the resistance of this process to extinction, suggests that targeting this psychological phenomenon can be a potential avenue for behavioral change to reduce the persistent motivational value of cues that leads to maladaptive pursuit as evidenced in substance use disorders (Robinson and Berridge, 1993; Marlatt, 1996; Conklin and Tiffany, 2002; Crombag et al., 2008; Zhang et al., 2009; Sinha, 2011).

By recording single neurons within the nucleus accumbens during occasion setting we discovered a widespread encoding of hierarchical stimuli that surpassed the number of neurons modulated by a simple Pavlovian conditioned stimulus. These data are reminiscent of the encoding of stimuli that indicate the availability of actions to be performed to earn reward (Ambroggi et al., 2008, 2011; Nicola, 2010; Richard et al., 2016; Sicre et al., 2020). In these scenarios, discriminative stimuli act to instruct when an action will directly lead to reward and resolve uncertainty surrounding the relationship between actions and reward, although the nature of training also results in these stimuli also being directly related to reward. Interestingly, neural activity and dopamine signaling in the nucleus accumbens core is also essential in these scenarios (Ambroggi et al., 2008, 2011; du Hoffmann and Nicola, 2016, 2016; Sicre et al., 2020). This is in contrast to relatively modest effects observed with lesions or inactivation of the nucleus accumbens core, and while dopamine release is necessary in the nucleus accumbens core for Pavlovian conditioned approach, the magnitude of effect is not as extreme as in these scenarios where stimuli disambiguate the relationship between cues and reward or actions and reward (Di Ciano et al., 2001; Blaiss and Janak, 2009; Chang et al., 2012; Saunders and Robinson, 2012; Chang and Holland, 2013; Fraser and Janak, 2017; Sicre et al., 2020). We observed that a subset of nucleus accumbens neurons exhibited context-specific encoding of the conditioned stimulus, switching their response from excitations to inhibitions at cue onset. This divergent encoding, and in particular the switch to inhibition when the conditioned stimulus was predictive of reward, is consistent with a disinhibitory circuit function for the nucleus accumbens to promote cue-triggered motivation (Taha and Fields, 2005, 2006; Krause et al., 2010; O’Connor et al., 2015; Yang et al., 2018; Lafferty et al., 2020; Vachez et al., 2021).

Here we reveal that a primary function of the mesolimbic dopamine system is to rapidly modulate the value attributed to reward-paired cues and, in turn, scale context-appropriate motivation. We suggest that this is a common behavioral and neurobiological process explaining the rapid modulation of behavior driven by changes in context or by internal states like hunger and thirst. This contrasts with the notion that mesolimbic dopamine release encodes the long-running value of cues via an extensive refinement of their value during reinforcement learning to drive behavior. That dopamine neurons can encode expected value does not imply that this encoding reflects their only function. Indeed, adaptive behavior requires the divergence of behavioral responding from the long-running value of environmental stimuli in the absence of new learning. This implies that the regulation of conditioned reward-seeking is the norm when it comes to the contribution of VTA dopamine neuron activity to reward-seeking. This context-dependent transformation of reward-associated cues into targets of desire by the mesolimbic dopamine system provides a novel framework for understanding the dysfunction of this system in psychiatric illness.

## METHODS

### Subjects

Subjects were 99 male and 18 female Long-Evans rats. For selective inhibition of dopamine neuron activity, TH-Cre rats expressing the bacterial recombinase Cre under the control of the tyrosine hydroxylase (TH) promoter, the rate-limiting enzyme for the synthesis of dopamine, maintained on a Long-Evans background were used (Witten et al., 2011). Rats were single-housed in ventilated cages with ad libitum access to food and water in a temperature- and humidity-controlled room and maintained on 12:12 light/dark cycle (lights on at 07:00). After recovery from surgery, feeding was restricted to maintain weights at ∼95% of ad libitum feeding weights. All behavioral procedures took place between 13:00 and 20:00. All procedures were approved by the Animal Care and Use Committees at Johns Hopkins University and the University of Minnesota and followed the recommended guidelines in the Guide for the Care and Use of Laboratory Animals: Eighth Edition, revised in 2011.

### Surgery

Rats were anesthetized with isoflurane (5% induction, 1-2.5% maintenance) and standard stereotaxic procedures were used. For optogenetic experiments, a Cre-dependent viral vector containing halorhodopsin (n=12; AAV5-Ef1a-DIO-eNpHR3.0-eYFP; titer 3.1 x 1012 viral particles/mL; University of North Carolina) or a control GFP virus (n=12; AAV5-hsyn-EGFP; titer 1 x 1013 viral particles/mL; University of North Carolina) was infused bilaterally into the VTA (AP: -5.8 ML: ±0.7 DV: -8.0 for males; DV -7.7 for females). A volume of 700 nL of virus was infused per site at a rate of 100 nL/min through a 31-G, gas-tight Hamilton syringe controlled by a Micro4 Ultra Microsyringe Pump 3 (World Precision Instruments). The needle was left in place for 10 minutes following the infusion to allow for diffusion of the virus from the injection site. In the same surgery, rats were bilaterally implanted with custom made optic fiber implants (300 μm glass diameter) targeted to the VTA (15° angle; AP: -5.8 ML: ±2.61 DV: -7.55 for males; DV -7.1 for females). For microinfusion experiments 22-gauge stainless steel cannula (Plastics One) were implanted 1 mm over the nucleus accumbens core (n=12; AP: +1.8 ML: ±1.5 DV: -6) or shell (n=12; AP: +1.6 ML: ±1 DV: -6.5). For electrophysiological recordings, a custom-built microdrive containing 16 50-μm tungsten wires soldered to 8-pin Omnetics connectors was lowered slowly to the nucleus accumbens core (n=3; AP: +1.8 ML: ±1.5 DV: -6.5) with a silver reference wire wrapped around a screw in contact with the cerebellum. For fiber photometric recordings (n=8), a volume of 800 nL of AAV5-CAG-dLight1.3b-GFP (titer 1.03 x 1013 viral particles/ mL; Addgene) was infused per site at a rate of 100 nL/ min through a 31-G, gas-tight Hamilton syringe controlled by a Micro4 Ultra Microsyringe Pump 3 (World Precision Instruments) into the nucleus accumbens core (AP: +1.3 ML: ±1.3 DV: -6.9 300nL, -6.7 500nL). The needle was left in place for 10 minutes following the infusion to allow for diffusion of the virus from the injection site. In the same surgery, rats were implanted with optic fiber implants (400 μm glass diameter) targeted just dorsal to the infusion site (AP: +1.3 ML: ±1.3 DV: -6.5). For all surgeries, implants were secured to the skull with 4-8 screws placed in the skull and dental acrylic. After surgery, rats received injections of cefazolin (70 mg/kg, subcutaneous) to prevent infection and carprofen (5 mg/ kg, subcutaneous) to relieve pain. Rats were allowed to recover for 10 days prior to the beginning of behavioral procedures, with at least 4 weeks passing before any optogenetic manipulations were performed.

### Apparatus

Behavioral testing occurred in Med Associates conditioning chambers (St. Albans, VT) housed in sound- and light-attenuating cabinets and controlled by a computer running MedPC IV software. In the center of one wall was a fluid receptacle located in a recessed port. On the opposite wall near the ceiling of the chamber was a white houselight (28 V) and to the right of the houselight was a white-noise generator (10-25 kHz, 20 dB). Outside of the behavioral chamber but within the sound-and light-attenuating cabinet was a red houselight (28 V) that provided background illumination during each behavioral session. Fluids were delivered to the port via tubing attached to a 60 mL syringe placed in a motorized pump located outside each cabinet.

### Occasion setting task

Training in the occasion setting task was identical to procedures in (Fraser and Janak, 2019). In brief, in a single session there were 3 trial types, 10 of each trial, with a 3.3 minute average intertrial interval. On reinforced trials, the illumination of a white houselight for 5 s, a 5 s gap, and the presentation of white noise for 5 s (OS-⟶CS trials) was followed by the activation of a reward pump containing 15% sucrose (w/v) for 5 s (∼0.18 mL of sucrose delivered). The other trials consisted of either the sole illumination of the houselight for 5s (OS Alone trials) or the presentation of the white noise for 5s (CS Alone trials). Behavioral equipment and responses were controlled by a computer running MedPC IV software (Med Associates). Across all experiments rats had 18 total days of training before any testing, recording, or manipulations.

### Occasion setter extinction

Following acquisition, a subset of rats underwent occasion setter extinction. In these sessions all trials were OS alone trials (30 trials per session, ITI as before). Following this, rats were tested for the effects of OS extinction in a session in which reward was withheld to solely assess the effects of this manipulation on what they had learned.

### Conditioned reinforcement

In these tests, following training we asked rats to learn a novel operant response to earn the cues previously used in the occasion setting task. Rats could either lever press or nosepoke at test to earn the brief 2 second presentation of the occasion setter, the conditioned stimulus, or the combination of these two cues (presented simultaneously to link the action directly to their joint earning without imposing timeouts in responding). These sessions were conducted in extinction and each test lasted 40 minutes.

### Testing for configuration of cues

To assess whether rats configured the occasion setter and conditioned stimulus, we conducted a test session in which probe trials presented the conditioned stimulus followed by the occasion setter. This was opposite to the arrangement used during training. We accomplished this by adding 5 of these additional CS⟶OS trials into the standard occasion setting preparation. We analyzed time spent in the port during the OS in this case as that would reflect configuration (i.e., the rats are combining the two cues into one stimulus and/or merely using the presentation of two cues to update their responding).

### Reward devaluation

In two sessions rats were tested for their ability to incorporate alterations in the value of the reward into their behavioral responding. To devalue the reward, we made use of sensory specific satiety. Prior to any testing rats were habituated to drinking the alternate reward, 15% maltodextrin, in their homecage. For each of the two tests rats had free access for one hour to either 15% sucrose (devalued condition) or 15% maltodextrin (valued condition). Weight and volume of the bottles containing these solutions was monitored before and after each session to monitor consumption. Immediately following, the rats were moved to the experimental chambers and tested in the occasion setting task but under extinction conditions (no reward was delivered).

### Optogenetic inhibition of ventral tegmental area dopamine neurons

For optogenetic experiments, rats were habituated to being connected via a ceramic mating sleeve bilaterally to 200 μm core patch cords (Doric), which were connected to a fiber optic rotary join (Doric), connected to a separate 200 μm patch cord that interfaced with a 532 nm DPSS laser (Opto-Engine LLC). Laser delivery was controlled by transistor-transistor logic pulses from MedPC SmartCTRL cards that interfaced with a Master9 Stimulus Controller (AMPI), which dictated the duration of stimulation. During tests, constant laser light (15-20 mW) was delivered bilaterally for a constant duration of 5.5s, beginning either before a trial, beginning 0.5 s before either the houselight or white noise (or the time in which they would be presented in CS Alone and OS alone trials), beginning 4.75 s into the houselight and terminating 0.25 s into the white noise, or for one test in 3 1.5 s second pulses separated by 0.5 s intervals during the houselight. The order of tests was randomly determined with 1-2 days of performance in the task without light delivery, but still tethered to patch cables, to allow for the mitigation of any possible observed carry-over effects at test. All tests were reinforced. The primary behavioral event of interest was the time rats spent in the port during the white noise stimulus.

### Local pharmacological manipulation of the nucleus accumbens

Rats were accustomed to the handling and infusion procedure for 1-2 days prior to infusion by being transported to the procedure room after a training session, dummy stylets removed, a flat cut 28-gauge injector inserted into each cannula, and dummy stylets replaced. On the day prior to testing, a regular injector extending 1 mm past the guide cannula was used to confirm cannula patency. At test rats received infusions of either saline, a mixture of the GABA-B and GABA-A agonists, baclofen and muscimol (1.0 mM and 0.1 mM, respectively), the dopamine receptor antagonist flupenthixol (100 mM), or a combination of the AMPA and NMDA receptor antagonists CNQX and MK-801 (20 mM each) infused in a volume of 0.3 μL over 1 minute. After 1 additional minute to allow for diffusion away from the infusion site, injectors were removed, dummy stylets were replaced, and rats returned to their homecage for 5-10 minutes before test. There were four tests for each rat, one in each condition, in a random order and with at least one day of retraining without manipulation between. Test sessions were reinforced.

### Fiber photometry of dopamine release

To assess dopamine responses to cue presentations in the NAc we measured dLight fluorescence using fiber photometry. On recording days rats were tethered to implants using a low autofluorescence cable sheathed in a stainless armored jacket (400 μM, 0.57 NA; Doric-Lenses). A fluorescence mini-cube (Doric-Lenses) transmitted light streams from 465 nm and 405 nm LEDs (Doric-Lenses), sinusoidally modulated at 211 Hz, and 330 Hz, respectively. LED power was set at ∼100 μW. Fluorescence from neurons at the fiber tip was transmitted back to the mini-cube, where it was passed through a GFP emission filter, amplified, and focused onto a high sensitivity photoreceiver (Newport, Model 2151). A real-time signal processor (RZ5P, Tucker Davis Technologies) running Synapse software modulated the output of each LED and recorded photometry signals, which were sampled from the photodetector at 6.1 kHz. Demodulation of the brightness produced by the 465-nm excitation, which stimulates dopamine-dependent dLight fluorescence, versus isosbestic 405-nm excitation, which stimulates dLight in a dopamine-independent manner, allowed for correction of bleaching and movement artifacts.

Rats were tested for responses to all cues prior to extinction of the occasion setter, probing for configural learning and responding after occasion setter extinction, and finally testing following extinction without reward delivery. One rat’s data was lost due to a computer error for just the recording prior to these manipulations.

For analysis, both signals were low-pass filtered (2 Hz), down-sampled to 40 Hz, and a least-squares linear fit was applied to the 405-nm signal, to align it to the 465-nm signal. This fitted 405-nm signal was used to normalize the 465-nm signal, where ΔF/F = (465-nm signal – fitted 405-nm signal)/fitted 405-nm signal. TTL inputs from MedPC were time stamped in the photometry data file.

### Electrophysiological Recordings

Electrical signals and behavioral events were collected from freely behaving animals with the OmniPlex (Plexon) recording system as in (Ottenheimer et al., 2018, 2020). Briefly, animals were connected via a 16-channel digital Omnetics headstage and data cable to a motorized commutator that interfaced with the OmniPlex system and recorded with a 40 kHz sampling rate. Behavioral events from MedPC hardware were directly timestamped into the OmniPlex data. Active wires were assessed in real-time and selected for recording based on manual inspection of waveforms across all 16 channels with the assistance of a Grass audio monitor. Units were initially pre-selected based on this manual inspection, but all channels and waveforms were subsequently sorted into units offline using Offline Sorter software (Plexon) following data collection, and any units that were not recorded throughout a behavioral session were discarded. Units were selected based on clustering among the first 3 components in PCA space, and further refined using waveform energy and slices of these features throughout the session. Cross-correlations were used to identify and discard contaminated units that were recorded on multiple channels. We only included neurons that were identified as single-units through careful examination of auto and cross correlations, plotting waveform features over time to ensure stability and continuity, and discarded units with more than 0.2% of spikes within a 2-ms refractory period. Following sorting in Offline Sorter all units were manually inspected in Neuroexplorer (Nex Technologies) for their response to relevant behavioral events to ensure fidelity throughout the session and lack of noise contamination. Following these steps data were pre-processed in Neuroexplorer to generate events of interest and exported to MATLAB for analysis. As we made use of a microdrive to lower the electrodes after each recording (in 160 μm increments) we ensured that active units that we recorded each day were unique by cross-referencing previous recordings on that wire for each session. Any channels that consistently had identical units were discarded.

### Histology

Following the conclusion of experiments, rats were deeply anesthetized with sodium pentobarbital and perfused transcardially with 4% paraformaldehyde. For rats with electrode implants, a brief 10 μA DC current was passed through each electrode to mark its final location in the brain. Brains were post-fixed for 24 hours in 4% paraformaldehyde, cryoprotected in 25% sucrose in 0.1 M NaPB, and then sectioned on a freezing cryostat at -20° C in 50 um sections. Brains with electrode or cannula implants were mounted onto Fisher SuperFrost Plus slides, dried, stained with cresyl violet solution (FD Neurotechnologies) and coverslipped with Permount mounting medium. For photometry and optogenetic experiments, sections were processed for detection of GFP and/or TH using fluorescent immunohistochemistry. Sections were first washed in 0.1 M PBS containing 0.2% Triton-X (PBST) for 20 minutes and blocked for 30 minutes in 0.1M PBST containing 10% normal donkey serum. Primary antibody incubation (mouse anti-GFP, 1:1500, Invitrogen A11120, Lot#2180270; rabbit anti-TH, 1:500, Fisher AB152M1, Lot # 3510772) occurred overnight at 4 C. The following day sections were washed in PBST for 15 minutes, blocked in PBS containing 2% normal donkey serum for 10 minutes, then secondary antibody incubation (Alexafluor 488 donkey anti-mouse, 1:200, Invitrogen A21202, Lot# 2229195; Alexafluor 594 donkey anti-rabbit, 1:200, Invitrogen A21207, Lot# 2145022) occurred at room temperature for 2 hours. Following 15 minutes of washing in PBS sections were mounted onto Fisher SuperFrost Plus slides and coverslipped with Vectashield mounting medium containing DAPI. Brain sections were imaged with a Zeiss Axio 2 microscope for the reconstruction of final placements of optic fibers and virus expression with the VTA or NAc.

### Statistics and Data Analysis

The primary behavioral metrics of interest for all tests was the time spent in port (indicated here as a percentage of the 5 s stimulus) during the white noise in the occasion setting task. We normalized behavioral responding by subtracting average time in port measured in a period 10 s prior to the presentation of any stimuli. We also computed differences between responding on trials in which the white noise would be reinforced (OS⟶CS trials) and trials in which the white noise (CS Alone) was presented alone or the houselight presented alone (OS Alone) to allow for an assessment of each individual rat’s representative ability to discriminate amongst the trial types.

### Data analysis and visualization were performed with

MATLAB (Mathworks) and Prism 9 (Graphpad). For optogenetic manipulations, repeated measure ANOVAs were used to compare time in port across trial types or difference scores between virus groups for each test separately. Two-way repeated measures ANOVAs were used to analyze the impact of microinfusions at test, with each drug separately being compared to data from the saline control session.

Photometry signals were z-scored to the 5s prior to the onset of the first event in each peri-event window. To assess cue responses, a 1s snippet was extracted from the z-scored peri-event signal for each cue and averaged to give a peak signal for each cue and trial type on each recording day and then averaged across pre-extinction and post-extinction recordings for each animal. Cue responses were quantified by calculating the maximum fluorescence and AUC from the average peak signal for each animal. For OS responses we made use of nonparametric tests, namely the Friedman statistic, to account for the structure of this data.

For electrophysiology data we first processed the activity of each individual neuron by counting spikes in 25ms bins spanning the session and smoothing with a half-normal filter (σ=5) over the preceding 25 bins. We then created PSTHs in 100ms around the onset of each trial (OS presentation, or lack thereof) using the smoothed traces. We averaged together the activity on OS alone, CS alone, and OS⟶CS trials to create each neuron’s PSTH for each trial type and then found the mean and standard error of these traces across groups of neurons. To detect neurons with responses to variables of interest, we adapted a kernel regression approach used previously (Ottenheimer et al., 2023). For each neuron, we predicted the activity on each trial (−11 to +12 s relative to CS onset) in 100ms bins using a series of kernels with windows that spanned multiple seconds consisting of 100ms bins. This approach allows detection of time-varying response patterns and the attribution of variance to co-occurring predictors. The predictors were as follows: OS onset (0 to 2 s relative to OS), OS offset (5 to 7 s relative to OS), CS onset (0 to 2 s relative to CS onset), CS offset (5 to 7 s relative to CS onset), OS⟶CS interaction (0 to 2 s relative to CS onset), and reward interaction (5 to 7 s relative to CS onset), as well as a single column predictor that increased with trial number. All predictors took on a value of one for the duration of their kernel, except for OS-⟶CS interaction and reward interaction, which were -1 on OS⟶CS trials and 1 on CS alone trials, allowing us to detect differences in these response patterns on the two trial types. We fit our model with glmnet with alpha = 0 (ridge penalty) and predicted held out trials in a 10-fold cross-validation. To find the significance of variables of interest, we fit reduced models where we removed their associated kernels: OS onset and offset to detect OS responses, CS onset and OS⟶CS interaction to detect CS responses, and OS⟶CS interaction to detect context modulation of CS responses. We found the F-test statistic for each variable of interest by comparing the variance explained by the full and reduced models, and then we generated a p-value by comparing the statistic to the distribution of F-test values generated on neural activity with trial onset times shuffled throughout the session. Neurons with p<0.05 were considered significantly modulated and plotting the activity of these neurons confirmed strong variable-specific modulation. Neurons with significant OS⟶CS interaction were called CS-context modulated (even if they had significant CS responses, too), the remaining CS-responsive neurons were called CS-general. All neurons with significant OS responses were called OS-responsive.

We only analyzed data from electrophysiology and fiber photometry data when rats exhibited greater responding, evidenced by time in port, during the CS on OS+CS trials relative to both CS alone and OS alone on the day of recording. Depending on the structure of the data, post-hoc tests were performed either with Bonferroni’s or Tukey’s method. For all statistical tests α=0.05.

## SUPPLEMENTARY MATERIAL RESULTS

### Representative training data from male and female rats and the rats in the optogenetic experiment

The occasion setting task is trained in phases, with progressive introduction of the different trial types. We present here data from male and female rats (n=18 per group) throughout this training (**Supplemental Figure 1A**). Both female and male rats acquire the task at a similar rate, but female rats tend to respond more than male rats (see **Figure 1**). In addition, we show that there were no differences in training performance for rats with either GFP or halorhodopsin expressed in the VTA (**Supplemental Figure 1B**).

### Manipulations of VTA dopamine activity outside of the occasion setter or conditioned stimulus fail to affect behavior

It has been demonstrated that pulsed inhibition of VTA dopamine neurons for many trials can result in conditioned inhibition, a phenomenon that is potentially related to occasion setting (Chang et al., 2018). We asked if delivery of 3, 1.5 s pulses of 532 nm light during the OS would recapitulate our behavioral effects. During this test there were no detectable differences between Halo and GFP rats in either reward-seeking (**Supplemental Figure 2B**; effect of group F_(1,22)_=3.414, p=0.0.781; effect of trial type F_(2,44)_=51.27, p<0.0001; interaction F_(2,44)_=4.032, p=0.0247; all p between GFP and Halo p>0.05, within group all p<0.001 relative to OS⟶CS trials) or discriminations (**Supplemental Figure 2C**; effect of group F_(1,22)_=0.3002, p=0.5893; effect of discrimination F_(1,22)_=1.797, p=0.1937). VTA dopamine neuron activity has been implicated in working memory and performance in this task requires the retention of the prior presentation of the OS to organize appropriate responding to the CS. To test a potential explanation of working memory, we delivered light for the final 250ms of the OS period through the first 250ms of the CS period (Cools and D’Esposito, 2011; Arnsten et al., 2012; Choi et al., 2020). Although we observed an overall effect of this manipulation on behavior for Halo rats relative to GFP control animals, there were no statistically significant differences between GFP and Halo rats for reward-seeking on any trial type, nor for did this affect within-group differences in total reward-seeking (**Supplemental Figure 2E**; effect of group F_(1,22)_=7.612, p=0.0115; effect of trial type F_(1,35)_=57.41, p<0.0001; interaction F_(2,44)_=0.7488, p=0.4788; all p between GFP and Halo p>0.05, no differences within group all p<0.001 relative to OS⟶CS) or discriminations (**Supplemental Figure 2F**; effect of group F_(1,22)_=0.01141, p=0.7387; effect of discrimination F_(1,22)_=0.9352, p=0.3440; interaction F_(1,22)_=0.9680, p=0.3359). Finally, we asked if light delivery selectively during reward delivery would alter behavior, given the prominent role of VTA dopamine neural activity at the time of reward in driving learning and updating behavior (Steinberg et al., 2013; Chang et al., 2016, 2017; Keiflin et al., 2019). Even with this manipulation, we observed no effect of light delivery on affecting behavior between GFP and Halo groups on overall reward-seeking (**Supplemental Figure 2H**; effect of group F_(1,22)_=4.002, p=0.0579; effect of trial type F_(1,33)_=34.5, p<0.0001; interaction F_(2,44)_=2.598, p=0.0858) or in the ability of individual rats to discern between reinforced and non-reinforced CS presentations (**Supplemental Figure 2I**; effect of group F_(1,22)_=0.2549, p=0.6187; effect of discrimination F_(1,22)_=1.327, p=0.2617).

### Neuropharmacological dissection of nucleus accumbens shell contributions to occasion setting

Dopamine antagonism within nucleus accumbens shell had an overall similar impact of reducing cue-triggered reward-seeking in the occasion setting task (**Supplemental Figure 3B**; effect of trial type F_(1,17)_=22.94, p<0.0001; effect of drug F_(1,11)_=165.4, p<0.0001; interaction F_(1,17)_=15.62, p=0.0002). Following flupenthixol administration reward-seeking was significantly lower on OS+CS (p<0.0001), CS alone (p=0.0007), and OS alone trials (p<0.0001) relative to saline, with no significant differences following flupenthixol between OS+CS trials and CS alone (p=0.8291) or OS alone (p=0.0667) trials. This was reflected by an overall impact of flupenthixol in reducing discriminations among trial types (**Supplemental Figure 3C**; effect of drug F_(1,11)_=51.51, p<0.0001; within each discrimination p<0.01). Dopamine antagonism in the nucleus accumbens shell significantly reduced intertrial port entries (**Supplemental Figure 3D**; t_11_=7.336, p<0.001) but not intertrial port time (**Supplemental Figure 3E**; t_11_=1.296, p=0.2215). As a result, functional dopamine signaling in the nucleus accumbens core and shell is equivalently necessary for the performance of occasion setting.

Inactivation of the nucleus accumbens shell reduced reward-seeking across all trial types (**Supplemental Figure 3F**; effect of trial type F_(1,15)_=38.22, p<0.0001; effect of drug F_(1,11)_=32.09, p=0.0001; interaction F_(1,21)_=5.369, p=0.0139) resulting in significantly reward-seeking relative to saline infusion on OS+CS (p=0.0011) and CS alone trials (p<0.0001) and eliminated differences in responding between OS+CS trials relative to OS alone trials (p=0.1514) but, interestingly, not CS alone trials (p=0.0388). Despite this, inactivation of the shell resulted in significantly reduced overall discrimination in the task (**Supplemental Figure 3G**; effect of drug F_(1,11)_=13.10, p=0.0040; within each discrimination p<0.05). In contrast to its effects on cue-triggered responses, inactivation of the nucleus accumbens shell did not have a significant impact on intertrial port entries (**Supplemental Figure 3I**; t_11_=1.697, p=0.1178) or intertrial port time (**Supplemental Figure 3H**; t_11_=0.4630, p=0.6524). Collectively these data indicate neural activity in both nucleus accumbens core and shell is necessary for occasion setting, with a more prominent impact of inactivation within the nucleus accumbens core.

Glutamate antagonism within the nucleus accumbens shell was without significant effect on reward-seeking (**Supplemental Figure 3J**; effect of trial type F_(1,14)_=44.39, p<0.0001; effect of drug F_(1,11)_=1.976, p=0.1875; interaction F_(1,15)_=0.1706, p=0.8315), discrimination among reinforced and non-reinforced trials (**Supplemental Figure 3K**; effect of drug F_(1,11)_=0.3291, p=0.5778), or intertrial port time (**Supplemental Figure 3L**; t_11_=0.2096, p=0.8378). Glutamate antagonism in the shell did however result in a significant increase in intertrial port entries (**Supplemental Figure 3M**; t_11_=3.548, p=0.0046).

**Supplemental Figure 1.**
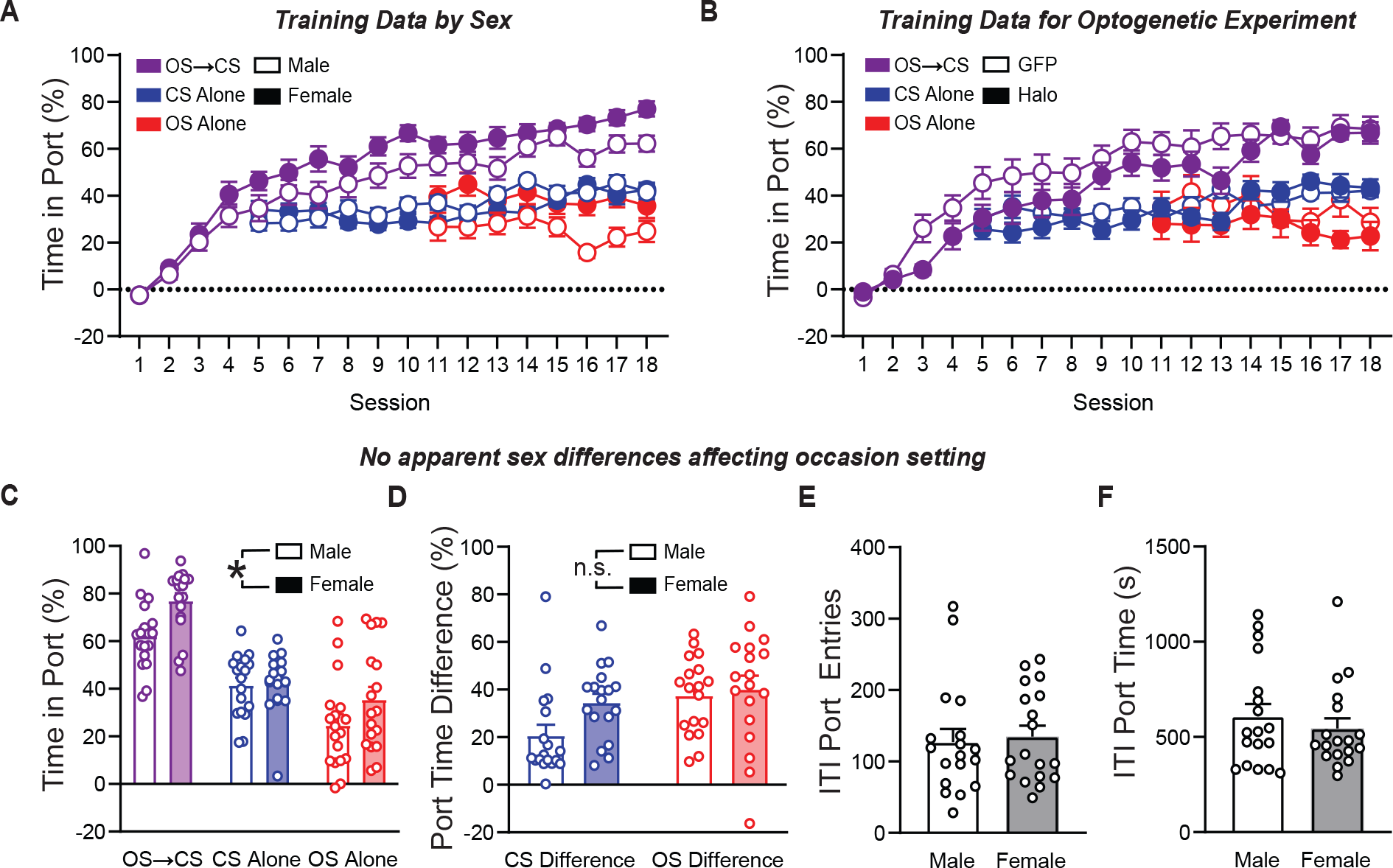
Representative training data in the occasion setting task. A. Conditioned responding as assessed by percent time in port for male and female rats during training (n=18 per group). B. Conditioned responding as assessed by percent time in port for TH-cre rats with either halorhodopsin or GFP expressed in VTA dopamine neurons (n=12 per group). C. Conditioned responding during occasion setting for male and female rats (n=18 each; effect of sex F(1,34)=6.416, p=0.0161; effect of trial type F(2,68)=66.17, p<0.0001; interaction of sex and trial type F(2,68)=2.021, p=0.1404). D. Difference in responding on OS⟶CS trials relative to CS alone and OS alone trials for male and female rats (effect of sex F(1,34)=4.075, p=0.0515; effect of discrimination F(1,34)=5.637, p=0.0234; interaction of sex and discrimination F(1,34)=1.384, p<0.2477). E. Intertrial port entries for male and female rats (t34=0.32, p=0.7509). F. Intertrial port time for male and female rats (t34=0.7491, p=0.4590). OS, occasion setter. CS, conditioned stimulus.

**Supplemental Figure 2.**
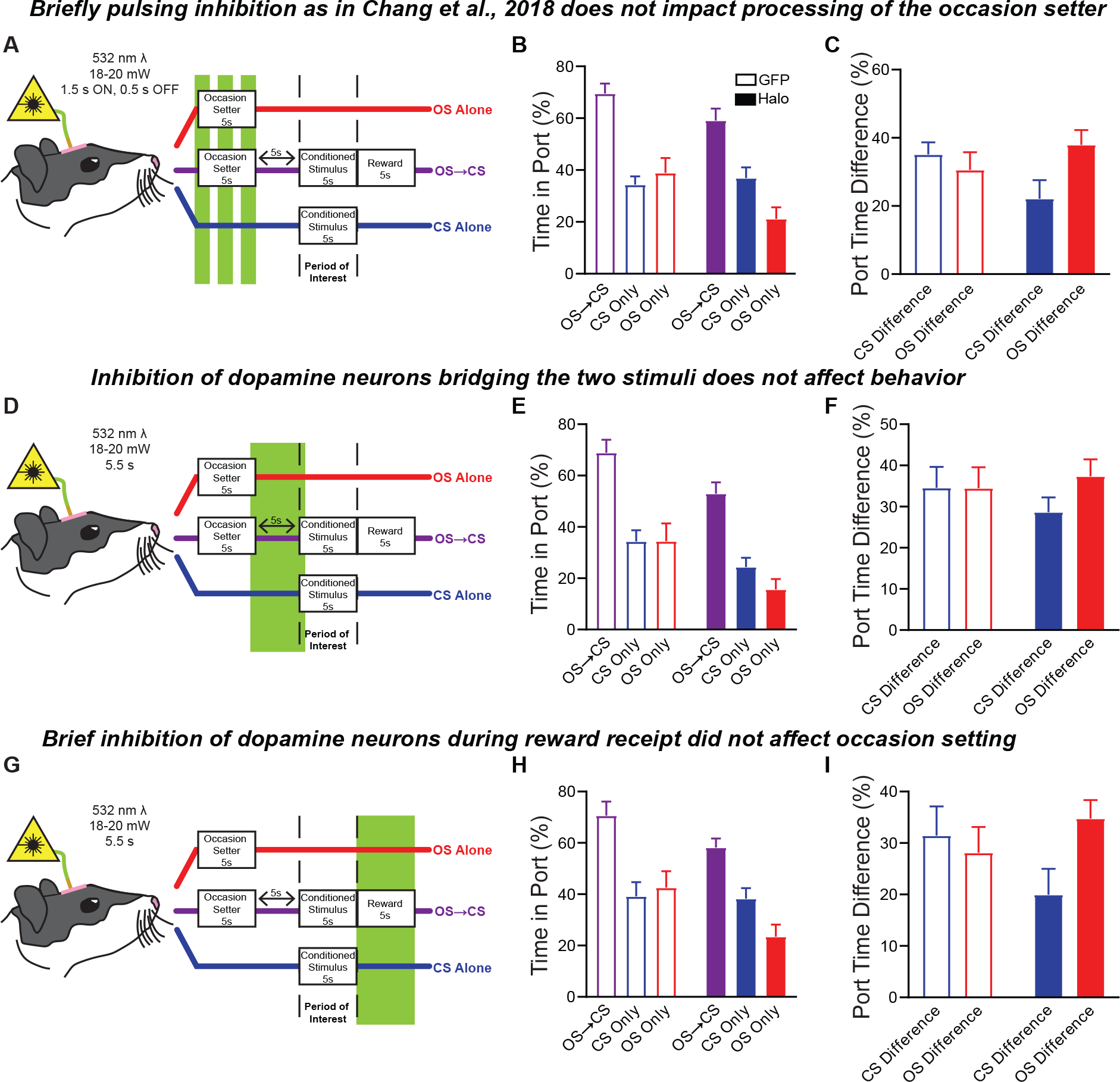
Inhibition of VTA dopamine neurons outside of the occasion setter or conditioned stimulus fail to affect behavior. A. Conditioned responding during occasion setting following 532 nm light delivery into the VTA for 3 1.5s, pulses during the occasion setter. E. Difference in responding on OS⟶CS trials relative to CS alone and OS alone trials following pulsed inhibition. F-G. Same as D-E but when light was delivered for 5.5s beginning 0.25s before occasion setter termination and ending 0.25s into the conditioned stimulus. H-I. Same as D-E but when light was delivered for 5.5s beginning 0.5s before the conditioned stimulus terminated. OS, occasion setter. CS, conditioned stimulus. eNpHr3.0, halorhodopsin. TH, tyrosine hydroxylase. ^*^p<0.05.

**Supplemental Figure 3.**
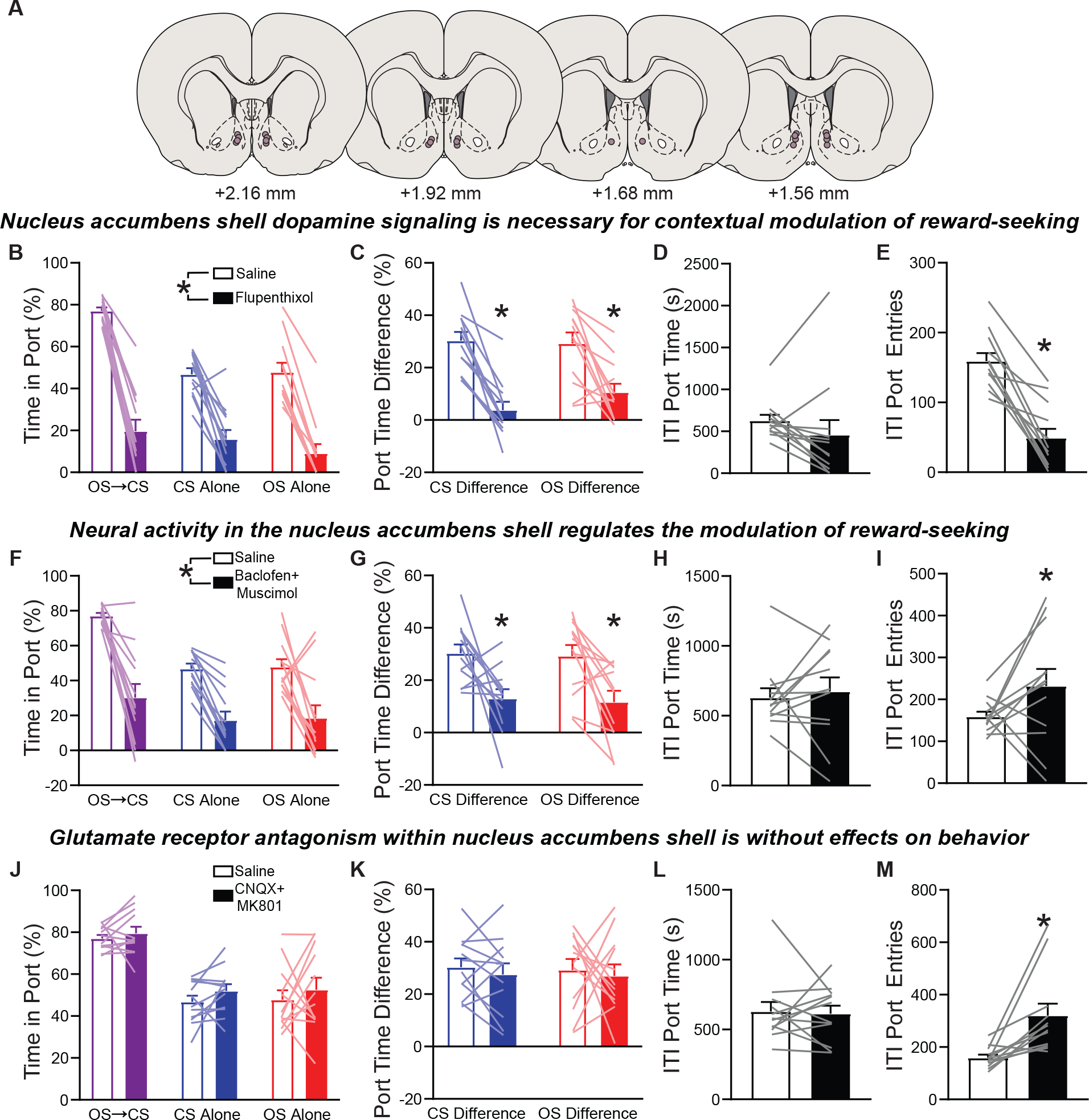
Nucleus accumbens shell dopamine and neural activity are essential for occasion setting. A. Recreation of microinjector tips within the nucleus accumbens shell (n=12). B. Conditioned responding as assessed by percent time in port following saline and flupenthixol infusions in the nucleus accumbens shell. C. Difference in responding on OS⟶CS trials relative to CS alone and OS alone trials following flupenthixol infusion. D. Intertrial port time following flupenthixol. E. Intertrial port entries following flupenthixol. F-I. Same as B-E but following infusion of baclofen and muscimol into the nucleus accumbens shell. J-M. Same as B-E but following infusion of CNQX and MK801 into the nucleus accumbens shell. OS, occasion setter. CS, conditioned stimulus. ^*^p<0.05.

